# *Vibrio cholerae* CsrA controls ToxR levels by increasing the stability and translation of *toxR* mRNA

**DOI:** 10.1101/2024.09.26.615275

**Authors:** Alexandra R. Mey, Charles R. Midgett, F. Jon Kull, Shelley M. Payne

**Author notes:** Corresponding author: Shelley M. Payne.

## Abstract

Regulation of colonization and virulence factor production in response to environmental cues is mediated through several regulatory factors in *Vibrio cholerae*, including the highly conserved RNA-binding global regulatory protein CsrA. We have shown previously that CsrA increases synthesis of the virulence-associated transcription factor ToxR in response to specific amino acids (NRES) and is required for the virulence of *V. cholerae* in the infant mouse model of cholera. In this study, we mapped the 5’ untranslated region (5’ UTR) of *toxR* and showed that CsrA can bind directly to an RNA sequence encompassing the 5’ UTR, indicating that the regulation of ToxR levels by CsrA is direct. Consistent with this observation, the 5’ UTR of *toxR* contains multiple putative CsrA binding sequences (GGA motifs), and mutating these motifs disrupted the CsrA-mediated increase in ToxR. Optimal binding of CsrA to a defined RNA oligonucleotide required the bridging of two GGA motifs within a single RNA strand. To determine the mechanism of CsrA regulation, we assayed *toxR* transcript levels, stability, and efficiency of translation. Both the amount of *toxR* mRNA in NRES and the stability of the *toxR* transcript were increased by CsrA. Using an in vitro translation assay, we further showed that synthesis of ToxR was greatly enhanced in the presence of purified CsrA, suggesting a direct role for CsrA in the translation of *toxR* mRNA. We propose a model in which CsrA binding to the 5’ UTR of the *toxR* transcript promotes ribosomal access while precluding interactions with RNA-degrading enzymes.

**IMPORTANCE:** *Vibrio cholerae* is uniquely adapted to life in marine environments as well as in the human intestinal tract. Global regulators such as CsrA, which help translate environmental cues into an appropriate cellular response, are critical for switching between these distinct environments. Understanding the pathways involved in relaying environmental signals is essential for understanding both the environmental persistence and the intestinal pathogenesis of this devastating human pathogen. In this study, we demonstrate that CsrA directly regulates synthesis of ToxR, a key virulence factor of *V. cholerae*. Under conditions favoring high levels of active CsrA in the cell, such as in the presence of particular amino acids, CsrA increases ToxR protein levels by binding to the *toxR* transcript and enhancing both its stability and translation. By responding to nutrient availability, CsrA is perfectly poised to activate the virulence gene regulatory cascade at the preferred site of colonization, the nutrient-rich small intestinal mucosa.

## INTRODUCTION

*Vibrio cholerae* is a Gram-negative bacterial pathogen associated with epidemics and pandemics of the disease cholera. Although most strains of *V. cholerae* are non-pathogenic inhabitants of marine environments, some *V. cholerae* are pathogenic and capable of transitioning from their native aquatic environments to the human host. Within the host, they colonize the small intestine and grow to large numbers before being shed back into the environment, thus completing the infectious cycle (reviewed in (1–3). Once pathogenic *V. cholerae* becomes endemic in natural waterways, future outbreaks are very difficult to prevent, especially in areas of poor sanitation resulting from natural, economic, or political disasters. Due to underreporting, it is difficult to assess the global cholera burden, but some studies estimate there are at least 2.9 million cases of cholera, and up to 95,000 associated deaths, per year worldwide (4).

*V. cholerae* pathogenesis depends on the production of several virulence factors, including the toxin co-regulated pilus (TCP) for attachment to, and colonization of, the small intestine (5), and the cholera toxin (CT), which causes the effusive watery diarrhea characteristic of the cholera disease (reviewed in (6, 7)). Virulence factor production in the intestinal environment relies on transcriptional regulators that coordinate the expression of virulence genes with the appropriate environmental cues, such as temperature, osmolarity, pH, oxygen status, and nutrient levels, as well as the presence of antimicrobial compounds such as bile, organic acids, and antimicrobial peptides (reviewed in (8–12).

One of the major virulence gene regulators in *V. cholerae* is ToxR, a cytoplasmic membrane-associated transcription factor that is required for both TCP and CT production and is essential for virulence (5, 13–15). ToxR, together with its integral membrane binding partner ToxS, and in conjunction with another transmembrane transcription factor, TcpP, regulates transcription of *toxT* (16–18). ToxT, in turn, induces transcription of the *ctxAB* genes, encoding the cholera toxin, and the *tcp* operon, encoding the TCP (reviewed in (2)). ToxR is also capable of directly activating *ctxAB* transcription but requires bile as a co-factor for this activity (19). In addition, ToxR controls the composition of the outer membrane by reciprocally regulating the levels of two major outer membrane porin (Omp) proteins, OmpT and OmpU: ToxR represses *ompT* transcription while activating *ompU* transcription (5, 20–22). OmpT is the more permissive of the two porins, and may be beneficial during growth in the aquatic environment, where nutrient concentrations are low; however, the more restrictive porin OmpU confers an advantage in the intestine, where inhibitory compounds such as antimicrobial peptides and bile acids are abundant (23, 24) (25). Thus, remodeling of the outer membrane represents a critical step in the transition from an aquatic habitat to the host environment.

How the ToxRS complex senses and responds to environmental cues is not completely understood. The activity of the ToxR protein can be directly altered by several environmental factors. Bile acids, which are present in the small intestine, greatly increase the activity of ToxR at the *omp* promoters, but without increasing the amount of *toxR* mRNA or ToxR protein (26). Similarly, the bacterial signaling molecule indole increases ToxR activity at the *leuO* promoter without altering *toxR* transcription or ToxR protein levels (27). Both bile and indole may to bind to the periplasmic domain of ToxR, but it is not known how the binding of these substances enhances the activity of ToxR at its target promoters (27–29).

Although the expression of *toxR* itself, both in terms of transcript levels (30) and protein production (31, 32), was not affected by growth of the bacteria in several laboratory conditions favoring virulence gene expression, leading to the hypothesis that *toxR* expression may be constitutive, studies in our laboratory have demonstrated that expression of the ToxR protein is not constitutive under all conditions but responds to specific components in the growth medium (26), adding another layer of complexity to the environmental regulation of the ToxR regulon. The ToxR level is low when cells are grown under nutrient-poor conditions, but increases rapidly if a mixture of specific amino acids, asparagine, arginine, glutamic acid and serine (NRES)(33), is added to the medium (26). The NRES-mediated increase in ToxR levels correlates with a switch in porin production from OmpT to OmpU (20, 26), consistent with elevated ToxR activity. These *in vitro* conditions may mimic the environmental shift experienced by *V. cholerae* during its transition from a nutrient-poor aquatic habitat to the nutrient-rich human small intestine.

Through further studies, we determined that the ToxR response to NRES is controlled by the posttranscriptional regulator CsrA (34). CsrA is a global regulator that controls cellular metabolism, physiology, and a number of pathways involved in virulence, including biofilm formation, motility, host-pathogen interactions, and virulence gene expression, in *V. cholerae* and other human pathogens (34–40). In *V. cholerae*, as in many other species (41–43), the *csrA* gene is essential, and a complete deletion of the *csrA* gene is not possible without deleterious effects on cell growth and viability; however, certain point mutations or C-terminal deletions resulting in partial CsrA activity are well tolerated (34, 38, 44). One such mutant, carrying a histidine residue in place of the arginine at position 6 in CsrA (*csrA*.R6H), had wild-type growth in culture, but did not exhibit the increase in ToxR and OmpU levels normally observed in response to NRES (34). The loss of ToxR regulation was not due to reduced levels of CsrA (45), and thus the R6H mutation may alter the affinity or specificity of RNA binding by CsrA. Not surprisingly, given the role of CsrA in regulating ToxR levels, the *csrA*.R6H mutant was severely attenuated in the infant mouse model of *V. cholerae* infection (34). *V. cholerae* CsrA was shown previously to affect expression of *V. cholerae* virulence factors through the quorum sensing regulator LuxO and the virulence gene regulator HapR (38), but no direct targets of CsrA regulation were identified in the LuxO-HapR pathway. In addition, we demonstrated that the NRES effect on ToxR levels is independent of both LuxO and HapR (34), suggesting that regulation of ToxR by CsrA is either direct, or involves a separate pathway from quorum sensing.

CsrA typically regulates its targets post-transcriptionally by binding to target RNA transcripts, often in the 5’ untranslated region (5’UTR), and either repressing or enhancing their translation or stability (reviewed in (36, 46)). CsrA recognizes a conserved motif in the RNA sequence, the hallmark of which is a conserved GGA motif located in the loop of an RNA stem-loop structure or in an otherwise unpaired region of the RNA. A common mechanism of negative regulation by CsrA is repression of translation initiation. This may involve binding of CsrA directly to the Shine-Dalgarno (SD) sequence, preventing the ribosome from loading, or CsrA promoting a particular folding of the RNA that sequesters the SD sequence and makes it inaccessible to the ribosome. CsrA binding may also promote mRNA degradation by facilitating access to the RNA by RNA degrading enzymes. Although less well studied, there are also examples of positive regulation by CsrA. One type of activation mechanism involves increasing the stability of an RNA structure in which the SD is maximally accessible for ribosome binding (47). Another example is the occlusion of access points for RNases when the transcript is bound by CsrA, thereby stabilizing the RNA (42). Our earlier studies concluded that the regulation of ToxR protein levels by CsrA in *V. cholerae* occurs via a positive regulatory mechanism (34), and in this study we show that CsrA binds to the 5’UTR of the *toxR* transcript and increases both the stability and translation of *toxR* mRNA.

## RESULTS

### *V. cholerae* CsrA is capable of binding directly to a sequence encompassing the 5’ untranslated region (5’UTR) of the *toxR* gene

*Delineation and characterization of the 5’ untranslated region (5’UTR) of* toxR. CsrA most commonly exerts its regulatory effects on protein synthesis by binding to the 5’UTR of the transcript. Figure 1A shows the putative promoter sequence, transcription start site (TSS)(+1), and 5’UTR of the *toxR* gene, but a definitive sequence for the *toxR* 5’UTR has not been published. The TSS for *toxR* was identified in two separate studies utilizing differential RNA-seq (dRNA-seq) to distinguish primary transcripts from processed transcripts by the Terminator 5’-Phosphate-Dependent Exonuclease (TEX) method (48, 49); however, two different translational starts of the ToxR protein have been proposed. The first in-frame ATG following the start of transcription (ATG1), positioned 72 nucleotides downstream of the transcription start site, was chosen as the most likely start of translation (14) based on the observed size of the ToxR protein in maxicell analyses (Miller, 1985; referenced in (14)). In contrast, the second in-frame methionine codon was considered to be the relevant start site (ATG2) based on N-terminal sequencing of the cytoplasmic domain of ToxR encoded on a plasmid expressing *toxR* from the *tac* promoter (50). To determine the true translational start of *V. cholerae* ToxR in N16961 during growth under the conditions used in these experiments, and therefore relevant for delineating the 5’UTR of *toxR* and understanding CsrA-mediated regulation, each of the two proposed ATG codons was mutated in the *V. cholerae* chromosome, and the effects of each mutation on ToxR production and activity were assessed. The ATG sequences were changed to CTT, which encodes leucine, a favorable amino acid substitution for methionine (51). The CTT codon has been shown in *E. coli* to lack any translation initiation activity (52).

**Figure 1.**
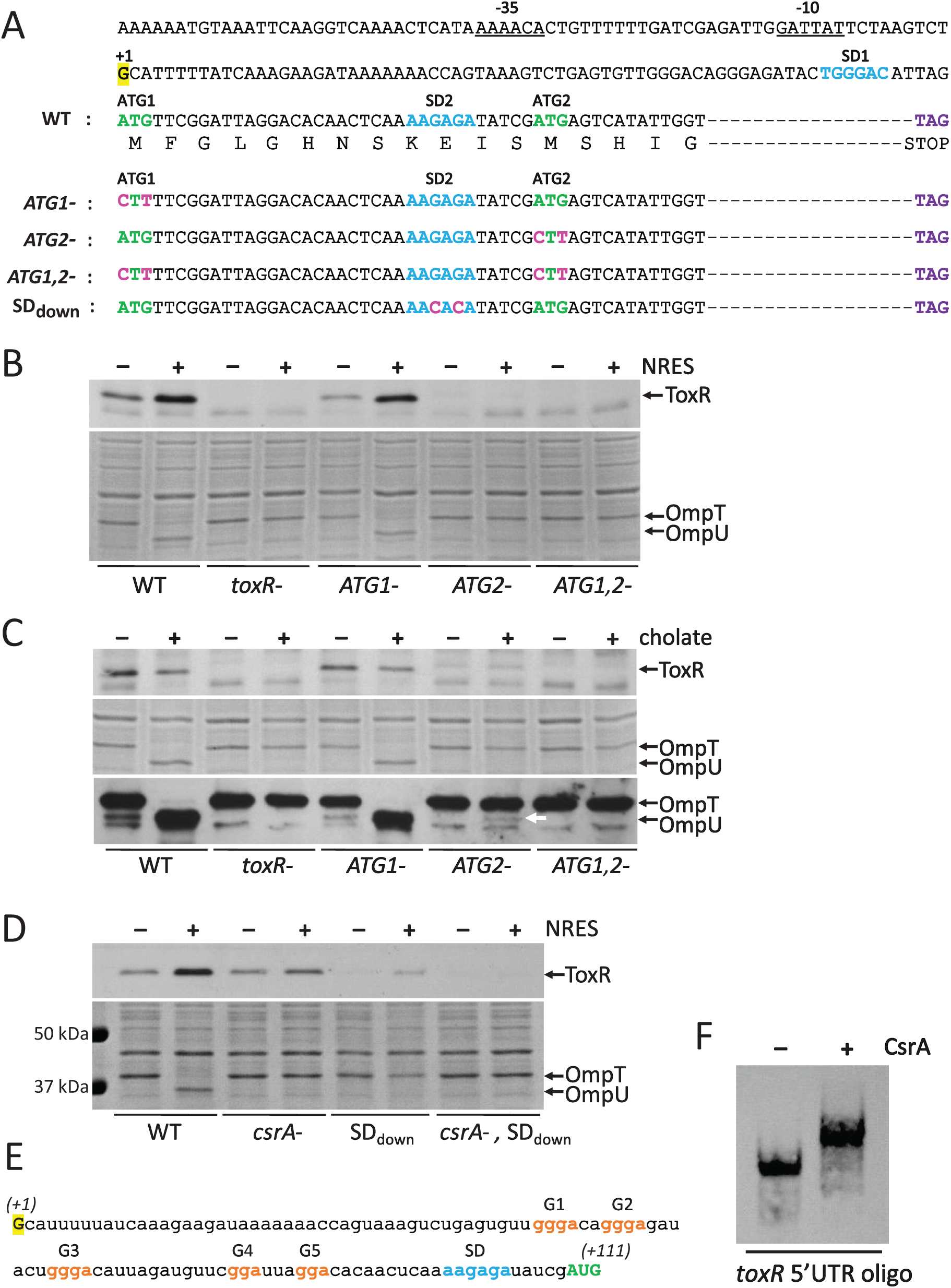
Identification and characterization of the 5’ untranslated region (5’UTR) of *toxR* and demonstration of CsrA binding to the 5’UTR sequence. (A-D) Delineation of the 5’UTR of *toxR* and identification of the start of translation and the associated Shine-Dalgarno (SD) sequence. (A) The promoter region and two potential start codons for translation of the *toxR* open reading frame are shown in the top part of the panel. The proposed -10 and -35 sequences for transcription initiation are underlined, and the start of transcription (+1), determined elsewhere by differential RNA-Seq (48, 49), is highlighted in yellow. The two candidate in-frame ATG codons (labeled ATG1 and ATG2) are shown in green downstream of their predicted SD sequences, which are shown in blue (SD1 and SD2, respectively). The encoded polypeptide sequence of the N-terminus of ToxR is indicated underneath the nucleotide sequence. The dashed line represents the part of the *toxR* coding sequence not shown. The stop codon is indicated in purple. The mutations introduced into the in-frame ATG codons and the SD2 sequence are shown in pink. (B) The wild-type strain N16961 (WT), the *toxR* mutant strain SAC119 (*toxR-*), the single mutants N*toxR*.M1L (*ATG1-*) and N*toxR*.M13L (*ATG2-*), and the double mutant N*toxR*.M1,13L (*ATG1-, ATG2-*) were grown to mid-logarithmic phase in T medium with or without 12.5 mM NRES mix. Whole cell preparations were resolved by SDS-PAGE and immunoblotted using polyclonal anti-ToxR antisera (top panel) or stained with Coomassie Blue to visualize the Omp proteins (bottom panel). (C) The strains referenced in (B) were grown to mid-logarithmic phase in T medium with or without sodium cholate. Whole cell preparations were resolved by SDS-PAGE and immunoblotted using polyclonal anti-ToxR antisera (top panel), stained with Coomassie Blue to visualize Omp proteins (middle panel), or immunoblotted using anti-Omp antisera to visualize both OmpT and OmpU. (D) The wild-type strain N16961 (WT), the *csrA* mutant strain N*csrA*.R6H (*csrA-*), and the SD2 mutant strains N*toxR*.SDdown (SDdown) and R6H.SDdown (*csrA-*, SDdown) were grown to mid-logarithmic phase in T medium with or without 12.5 mM NRES mix. Whole cell preparations were resolved by SDS-PAGE and immunoblotted using polyclonal anti-ToxR antisera (top panel) or stained with Coomassie Blue to visualize the Omp proteins (bottom panel). (E) The RNA sequence of the 5’UTR and translation initiation codon of *toxR* based on mapping of the translational start and SD sequence for the *toxR* gene above. (F) *V. cholerae* CsrA binds to the 5’UTR of the *toxR* transcript. An RNA electrophoretic mobility shift assay (REMSA) was performed to demonstrate direct binding of CsrA to an RNA oligonucleotide representing the entire 5’UTR and initiation codon of *toxR*, starting at the transcriptional +1 nucleotide and ending after the second in-frame AUG codon (111 nucleotides), as shown in (E). The REMSA binding reactions with the *toxR* 5’UTR oligonucleotide and purified *V. cholerae* CsrA were resolved in a non-denaturing 10% polyacrylamide gel, blotted to a positively charged membrane, and developed as described in the Methods section to visualize the position of the biotinylated RNA probe.

To discriminate between the two start codons, we first determined ToxR protein levels in the mutants grown in the presence or absence of NRES. Mutating ATG1 did not significantly affect the amount of ToxR produced, either with or without NRES (Fig. 1B, upper panel). Like the wild-type parental strain, the ATG1 mutant strain produced increased ToxR in the presence of NRES. This shows that translation from ATG1 is not essential for ToxR production under these conditions, and it suggests that the second in-frame ATG is active. When the second ATG was mutated, the ToxR protein was virtually undetectable, whether or not NRES was present, showing that ATG2 is the major initiation codon for *toxR* translation initiation (Fig. 1B). Upon longer exposure, a faint band became visible in the ATG2 mutant strain that was not present in the *toxR* null strain, indicating that a low amount of ToxR was being synthesized (Fig. S1). The weaker band appeared to be running a little higher than the dominant ToxR band, consistent with the ATG1-derived ToxR protein being 12 amino acids longer than the major ToxR protein species (Fig. S1). This suggests that, although ATG2 is the main translation initiation codon, a very low level of translation initiation takes place from ATG1. The ToxR protein translated from ATG1 did not appear to be regulated in response to NRES (Fig. S1). To confirm that the ToxR protein detected in the ATG2 mutant strain is translated from ATG1, and not from the mutated codon (CTT) or from another cryptic start codon in the *toxR* 5’UTR, a double ATG mutant was created and tested for ToxR production. The weak ToxR band present in the ATG2 mutant strain was not detected in the double ATG mutant (Fig. 1B, upper panel, and Fig. S1), showing that this, much less abundant, ToxR species is indeed translated from ATG1.

To verify that the observed levels of ToxR protein in the ATG mutants could be correlated with ToxR transcriptional activity, we analyzed the outer membrane protein profiles of the mutant strains (Fig. 1B, lower panel). ToxR activates *ompU* and represses *ompT* expression, and a change in the level or activity of ToxR can be easily detected by visualizing the Omp profiles in Coomassie-stained gels (26, 34). The ATG1 mutant exhibited an outer membrane protein pattern indistinguishable from that of the parental wild-type strain, with high levels of OmpU in response to NRES, as expected with the ATG1 mutant strain producing wild-type levels of ToxR and having a normal NRES response (Fig. 1B, lower panel). The ATG2 mutant, however, did not undergo the switch from OmpT to OmpU in response to NRES (Fig. 1B, lower panel), in agreement with the low amounts of ToxR produced in the absence of ATG2. As expected, no OmpU was detected in the double ATG mutant.

We have shown in earlier work that the ToxR protein can be activated by exposure to bile acids, including cholate and deoxycholate (26), even when the basal amount of ToxR protein is very low, such as during growth in minimal medium or in the *csrA* mutant (Fig. S2). Therefore, the cholate assay is a very sensitive assay for detecting ToxR function when the level of ToxR is below the normal threshold for activity. To ascertain whether the longer ToxR variant detected in small amounts in the ATG2 mutant strain has any activity, the response to cholate was analyzed. As observed previously, cholate did not increase ToxR production (Fig. 1C, upper panel), but a low level of activity of the variant protein was observed. Although OmpU was not visible in the stained gel of whole cell lysates of the ATG2 mutant strain grown in the presence of cholate (Fig. 1C, middle panel), a faint band corresponding to OmpU was detected by Western blot analysis (Fig. 1C, lower panel), indicating that ToxR protein produced from ATG1 can be activated by cholate to promote *ompU* expression, albeit at a very low level, consistent with the low amounts of ToxR made in this strain (Fig. 1C, upper panel). As expected, no OmpU was detected in either the stained gel or by Western blotting in the double ATG mutant, confirming the total absence of ToxR in this strain. Taken together, this demonstrates that the primary initiation codon for *toxR* translation is the second in-frame ATG, but a small amount of ToxR with low activity can be produced from the first in-frame ATG following the start of transcription.

#### Significance of the open reading frame between the two in-frame ATGs

A study using a ToxR-responsive *ctx-lacZ* reporter assay in *E. coli* to determine ToxR activity found that a ToxR variant initiating at the second in-frame ATG was inactive compared with ToxR initiating at the first ATG (53), suggesting that the sequence between the two ATGs may be necessary for ToxR function. To test whether the open reading frame between the two in-frame ATG codons is relevant for ToxR production or activity, two different frame-shift mutations were introduced immediately following the first ATG. Insertion of a thymine base in this position (FS1) (Fig. S3A) created a new open reading frame in the sequence between the two in-frame ATGs. When this strain was grown in minimal medium with and without NRES supplementation, ToxR protein was made, and the level of ToxR was responsive to NRES (Fig. S3B upper panel). Consistent with this result, OmpU was made in response to NRES in the FS1 mutant strain (Fig. S3B, lower panel). Because robust amounts of ToxR and OmpU protein were made in the FS1 mutant, these results indicate that the signals for translation from ATG2 were not significantly affected by the early frame-shift, and the sequence of amino acids between the two ATG codons is likely not critical for either ToxR production or ToxR function. Deletion of the T immediately following the ATG1 codon (FS2) caused a premature stop immediately following the ATG (Fig. S3A). This mutant also displayed a relatively normal NRES response, with an increase in ToxR (Fig. S3B, upper panel) and a switch from OmpT to OmpU (Fig. S3B, lower panel), showing that translation of the intervening sequence between the two in-frame ATGs is not necessary for producing active ToxR from the second ATG. As an additional test of ToxR function, the FS1 and FS2 mutants were grown in the presence of cholate; both strains exclusively produced OmpU in response to cholate, showing that the ToxR protein made in each of the mutants with a frame-shift between ATG1 and ATG2 is responsive to bile activation and fully functional in terms of regulating *ompU* and *ompT* expression (Fig. S3C).

#### Identification of the Shine-Dalgarno (SD) sequence for toxR translation

CsrA is known to exert regulatory effects on translation, often by freeing up or occluding the Shine-Dalgarno (SD) sequence for ribosome binding. To identify the SD sequence used for translation of the *toxR* transcript, mutational analysis of the predicted SD sequence upstream of the second ATG (SD2) was carried out. Two transversion mutations were introduced, changing the two conserved G nucleotides to C nucleotides (Fig 1A). The resulting mutant, SDdown, had significantly lower ToxR levels, demonstrating the importance of the proposed SD sequence for *toxR* translation, further establishing the second in-frame ATG as the true start of ToxR (Fig. 1D, upper panel). Interestingly, the SDdown mutant still responded to NRES and CsrA, producing the most ToxR protein when grown in the presence of NRES, but only when CsrA was intact, suggesting that the sequence of the SD itself may not be directly involved in the regulation by CsrA. The amount of ToxR made in the SDdown mutant strains was not high enough to induce OmpU synthesis in response to NRES (Fig. 1D, lower panel); however, a small amount of OmpU was made in the SDdown mutants in the presence of cholate, indicating the presence of bile-activated ToxR, albeit at a much lower level than in the wild-type strain, as the switch from OmpT to OmpU was incomplete (Fig. S4A). It is possible that the mutations introduced into the SD did not completely eliminate ribosome binding, or another sequence may serve as a site for ribosome loading.

#### V. cholerae CsrA binds to the toxR 5’UTR

With the identification the main translation initiation codon for ToxR synthesis, we have delineated the *toxR* 5’UTR as the 108 nucleotides between the start of transcription (+1) to the start of translation from ATG2 (Fig. 1A). To determine if *V. cholerae* CsrA is capable of binding directly to the *toxR* 5’UTR, an RNA electrophoretic mobility shift assay (REMSA) was performed using a 5’ biotinylated RNA oligonucleotide comprising the entire 5’UTR and initiation codon of *toxR*, starting at the transcriptional +1 nucleotide and ending after the second in-frame AUG codon (111 nucleotides)(Fig. 1E). Electrophoresis of the biotinylated probe, pre-incubated with or without purified CsrA, was carried out to determine binding of CsrA to the RNA oligonucleotide. A clear retardation in the migration of the oligonucleotide in the presence of CsrA was detected (Fig. 1F), signaling that CsrA is capable of binding directly to the sequence encompassing the 5’UTR of *toxR*.

### CsrA is not required to disrupt the predicted SD-sequestering hairpin or to overcome limitations of a potentially weak SD sequence

The direct binding of CsrA to the 5’UTR of the *toxR* transcript has implications for how CsrA regulates ToxR production. In silico structure predictions for the folding of the 3’ half of the *toxR* 5’UTR suggested that the SD sequence may pair with nearby bases to form a stem structure that could sequester the SD and make it inaccessible to the ribosome (Fig. 2A). This prompted the hypothesis that CsrA binding to the *toxR* 5’UTR may prevent the formation of this stem structure, thereby promoting ribosome access to the SD sequence and increasing the efficiency of *toxR* translation. Two of the nucleotides predicted to pair with the SD sequence were mutated to eliminate the potential stem structure. The resulting disruption of the stem in the predicted secondary structure was supported by in silico analysis (Fig. 2A). The SDstem mutant was then evaluated for its dependence on NRES and CsrA for ToxR production. The results showed that the level of ToxR remained low in the SDstem mutant in T medium without NRES, and increased in the presence of NRES, just like in the wild-type parental strain (Fig. 2B, upper panel). Very little ToxR was made in the *csrA*, SDstem double mutant (Fig 2B, upper panel), showing that the SDstem mutant still requires CsrA for optimal ToxR production. The SDstem mutants each exhibited Omp switching in response to NRES (Fig. 2B, lower panel) and cholate (Fig. S4B) comparable to that of their parental strain, signaling that ToxR expression and function was not affected by the SDstem mutations. Thus, disrupting the specific hairpin that may be sequestering the SD sequence did not circumvent the need for NRES or CsrA in increasing ToxR levels; however, we cannot rule out that the overall folding of the 5’UTR in the absence of CsrA causes the SD sequence to be inaccessible.

**Figure 2.**
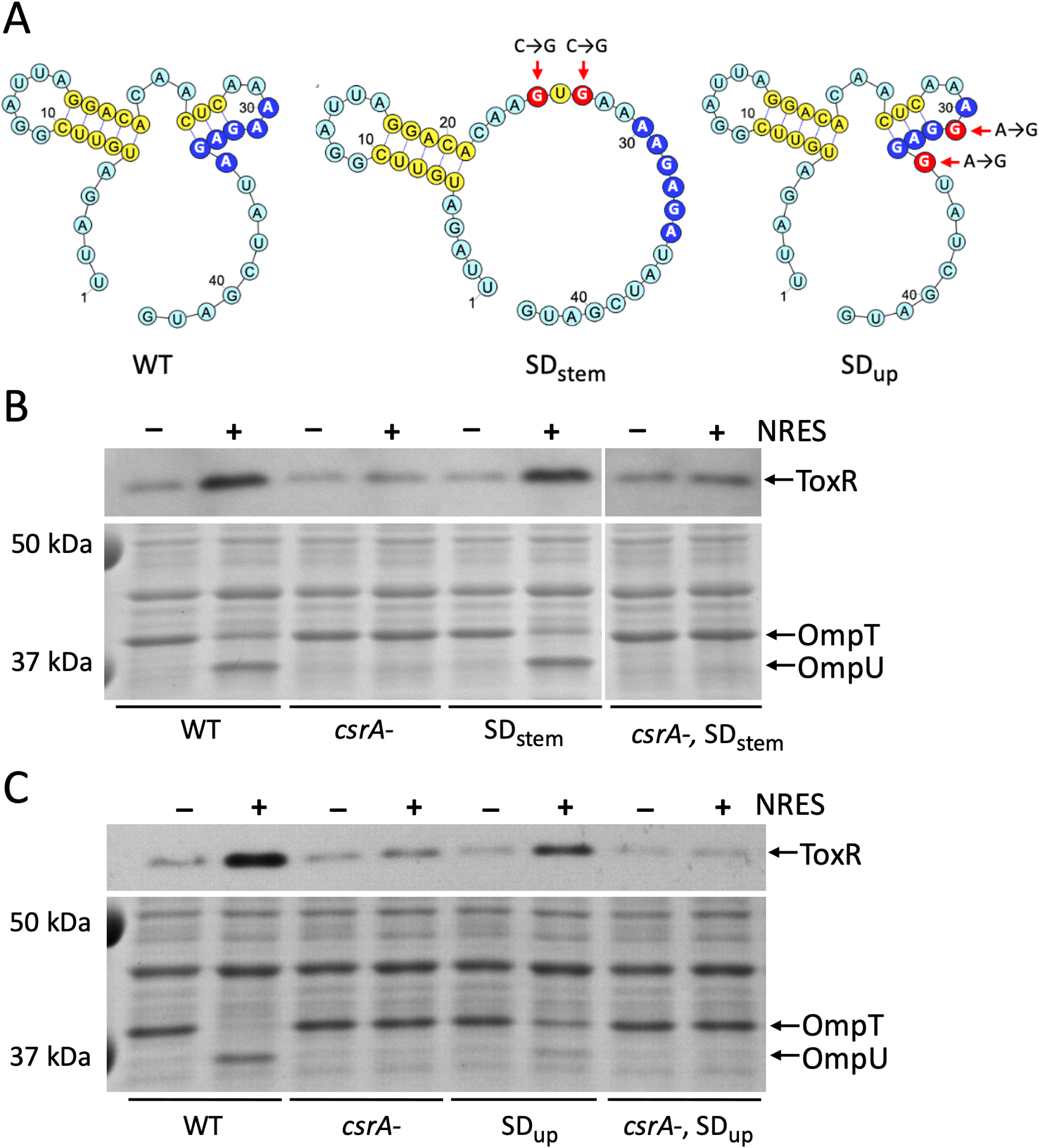
Evaluation of mutations that may eliminate the need for CsrA by increasing ribosome access and translation of *toxR*. (A) In silico folding of the wild-type (WT) sequence surrounding the *toxR* SD (dark blue circles) shows the formation of a potential stem structure that may sequester the SD and prevent ribosome access. The two mutations (red circles) predicted in silico to disrupt the putative stem and release the SD sequence are indicated with red arrows in the SDstem diagram. Mutations proposed to strengthen the SD sequence (red circles in the SDup diagram) did not alter the predicted folding of the region compared with the WT sequence. (B) To test whether eliminating the predicted stem structure alleviates the requirement for NRES or CsrA, the wild-type strain N16961 (WT), the *csrA* mutant N*csrA*.R6H (*csrA-*), and the SDstem derivative mutants N*toxR*.SDstem (SDstem) and R6H.SDstem (*csrA-*, SDstem) were grown to mid-logarithmic phase in T medium with or without 12.5 mM NRES mix. Whole cell preparations were resolved by SDS-PAGE and immunoblotted using polyclonal anti-ToxR antisera (top panel) or stained with Coomassie Blue to visualize the Omp proteins (bottom panel). The white vertical lines indicate that intervening lanes containing duplicate samples were removed from the gel for clarity. (C) To test whether altering the SD sequence to more closely match the SD consensus sequence can alleviate the requirement for NRES or CsrA, the wild-type strain N16961 (WT), the *csrA* mutant N*csrA*.R6H (*csrA-*), and the SDup derivative mutants N*toxR*.SDup (SDup) and R6H.SDup (*csrA-*, SDup) were grown to mid-logarithmic phase in T medium with or without 12.5 mM NRES mix. Whole cell preparations were resolved by SDS-PAGE and immunoblotted using polyclonal anti-ToxR antisera (top panel) or stained with Coomassie Blue to visualize the Omp proteins (bottom panel).

The SD sequence identified upstream of the second in-frame ATG (AAGAGA) is not a complete match with the consensus SD from *E. coli* (AGGAGG)(Shine and Dalgarno, 1974). It is possible that the genome-wide consensus SD sequence in *V. cholerae* does not match that of *E. coli*; however, a search of the *V. cholerae* N16961 genome (as described in Materials and Methods) revealed 8 putative 16S ribosomal RNA genes, all of which carry anti-SD sequences at the 3’ end that match 100% with the anti-SD of the *E. coli* 16S rRNA gene *rrsA*. This suggests that the basis for ribosomal recognition of the mRNA initiation sequences is similar in both species. Any deviation from the SD consensus may require other factors to increase ribosome recognition and translation initiation. Thus, the role of CsrA in *toxR* translation could be to increase ribosome loading at a relatively weak SD sequence. To determine whether the strength of the *toxR* SD sequence is a determining factor in CsrA regulation of ToxR production, the *toxR* SD sequence (AAGAGA) was mutated in both the wild-type and the *csrA* mutant background to match the consensus for a strong SD sequence (AGGAGG; SDup)(Fig. 2A). The resulting strains were tested for their ability to produce ToxR in the absence of NRES or CsrA. Rather than increase ToxR, the mutations decreased overall ToxR levels (Fig 2C, upper panel). Further, stimulation of ToxR production still required both NRES and CsrA, showing that the NRES and CsrA requirement in *toxR* translation was not alleviated by increasing the strength of the SD sequence (Fig. 2C, upper panel). Omp switching was less efficient in the strains carrying the SDup sequence (Fig. 2C, lower panel) as expected based on the slightly lower ToxR levels in these strains; however, the SDup mutants made a complete switch from OmpT to OmpU in the presence of cholate, indicating that the level of ToxR produced was still above the threshold for activation by bile (Fig. S4C). These results indicate that the role of CsrA is not simply to overcome the effects of a weaker SD sequence on translation efficiency.

### *V. cholerae* CsrA requires two adjacent RNA stem-loop structures with GGA motifs for efficient binding

CsrA is known to bind to a conserved sequence which includes an invariable GGA motif (GGGA in some targets) typically located within the loop of an RNA stem-loop structure or in an otherwise unpaired stretch of RNA sequence (46, 54). From studies in *E. coli* (55) and other species (56, 57), it is known that CsrA is active as a homodimer, and preliminary structural analyses of *V. cholerae* CsrA have confirmed that *V. cholerae* CsrA also forms homodimers in vitro (C. Midgett, manuscript in preparation). The presence of multiple putative CsrA binding sites in the *toxR* 5’UTR suggested that the *V. cholerae* CsrA dimer may bind more than one site at a time. In order to better understand the interactions of *V. cholerae* CsrA with its target transcripts, we undertook a study of the minimum binding requirements for CsrA-RNA interactions in *V. cholerae*. Among the best-known targets of CsrA binding in *V. cholerae* are the CsrB, CsrC, and CsrD sRNAs (38, 45), homologs of the CsrB and CsrC sRNAs in *E. coli*, which downregulate CsrA activity by binding and sequestering CsrA (54, 58).

Structural predictions of the *V. cholerae* sRNAs show numerous stem-loop structures with GGA (in some cases GGGA) motifs located in loops (38, 45), similar to the *E. coli* Csr sRNA structure predictions (54, 58). To study the minimal requirements for *V. cholerae* CsrA-RNA binding, we used short RNA oligonucleotides derived from CsrB as the substrate in REMSAs with purified *V. cholerae* CsrA. We first designed an oligonucleotide representing a short sequence of CsrB predicted by mfold (59) to fold into two adjacent stem-loop structures with a GGA motif in each loop (Fig. 3A), one of which is the expanded GGGA motif often found in CsrB homologs (38, 54). Of note, three of the five potential CsrA-binding motifs found in the *toxR* 5’ UTR have this expanded GGGA sequence (Fig. 1E). Electrophoresis of the biotinylated probe, pre-incubated with CsrA, showed a significant shift in migration of the oligonucleotide in the presence of increasing concentrations of CsrA (Fig. 3B), demonstrating binding of CsrA to the CsrB oligonucleotide. To test the role of the GGA motifs in CsrA binding, oligonucleotides were generated containing mutations in one or both of the GGA motifs in the CsrB oligo. CsrA failed to bind to oligonucleotides in which either or both of the (G)GGA motifs had been mutated (Fig. 3C), even though the predicted structures of the oligonucleotides remained unchanged by mfold predictions (Fig. S5). This confirms that *V. cholerae* CsrA recognizes (G)GGA motifs in the target RNA and suggests that CsrA requires two intact adjacent GGA motifs for efficient binding, even when the predicted overall structure of the oligonucleotide remains intact (Fig. S5). The small amount of the mut-right oligonucleotide that was shifted in the presence of CsrA (Fig. 3C) suggests it may be possible for CsrA to bind very weakly to an RNA containing only a single GGA motif; however, efficient binding clearly requires both GGA sequences. Further, RNA oligonucleotides representing only one of the stem-loop structures (CsrB-left and CsrB-right), each with only a single GGA motif, did not bind to CsrA when used either by themselves or together in the binding reaction (Fig. 3D). This confirms that a single stem-loop structure with a GGA motif does not a provide a good ligand for binding, and that the two GGA motifs in an RNA target must be physically linked in order for CsrA to bind effectively. Taken together, these observations suggest that optimal CsrA binding requires the bridging of two GGA binding sites within a single RNA strand.

**Figure 3.**
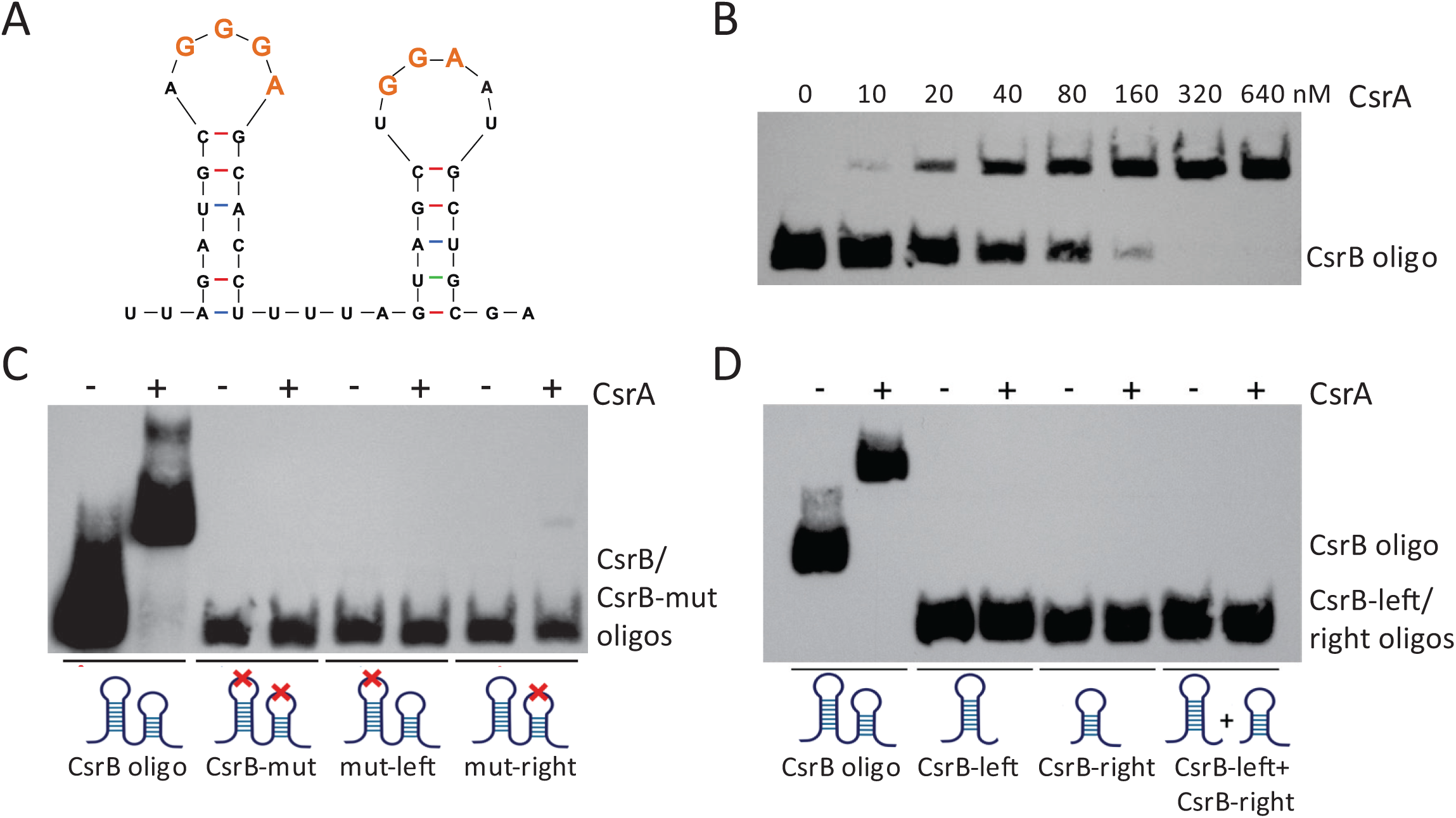
CsrA requires two GGA motifs within the same RNA strand for optimal binding. (A) Structure of an in silico-folded partial CsrB sequence (CsrB oligo) showing two adjacent stem-loop structures with (G)GGA motifs located in the loops. The structure folded structure was generated by mfold (59). (B) The CsrB oligo (0.1 nM) was mixed with increasing concentrations (0-640 nM) of purified *V. cholerae* CsrA, and RNA electrophoretic mobility shift assays (REMSAs) were performed as described in the legend to Figure 4. (C) REMSAs were performed with 1 nM each of CsrB oligo, oligo-mut (mutations in both (G)GGA motifs), mut-left (mutations in left loop GGGA motif), and mut-right (mutations in right loop GGA motif), and 640 nM purified *V. cholerae* CsrA, as described in the legend to Figure 1. The positions of the mutations in the predicted oligonucleotide structures are indicated with a red x in the cartoons below the image of the developed blot. (D) REMSAs were performed with 0.1 nM of the CsrB oligo, and 1 nM each of oligos CsrB-left (the left stem-loop of CsrB oligo), CsrB-right (the right stem-loop of CsrB oligo), or CsrB-left + CsrB-right, and 640 nM purified *V. cholerae* CsrA. The predicted structures of the oligonucleotides are represented by the cartoons below the image of the developed blot. The folding of each of the indicated oligonucleotides was predicted by mfold (59).

### Role of the conserved GGA motifs in the *toxR* 5’UTR

The *toxR* 5’UTR sequence contains 5 (G)GGA sequences, consistent with the observed binding of CsrA directly to this sequence. The (G)GGA motifs are located in close proximity to each other, spanning the region +47 through +87 from the start of transcription. The motifs are numbered 1-5, according to their order (5’ to 3’) in the sequence (Fig. 4A). RNA folding predictions for the *toxR* 5’UTR sequence shows that motifs 1, 2, and 4 are predicted to be located in unpaired loop regions, while motifs 3 and 5 are sequestered within paired stem structures (Fig. 4A). To determine the significance of the (G)GGA motifs for CsrA regulation of ToxR synthesis, a series of mutations in the *toxR* 5’UTR GGA motifs were introduced into the chromosome of *V. cholerae* (Fig. 4B), and the phenotypes of the mutants were determined. Mutations were designed to preserve as much of the predicted overall folding of the 5’UTR as possible, as determined by in-silico comparisons of predicted mutated and wild-type 5’UTR structures. Mutating G1, G2, and G3 significantly reduced the effect of NRES on ToxR production, and this strain produced a near-basal level of ToxR in the presence of NRES (Fig. 4C, top panel). Further, OmpU was not visible by Coomassie staining in this strain grown in the presence of NRES (Fig. 4C, bottom panel), consistent with the low ToxR level. This suggests that one or more of these 3 motifs plays a critical role in the regulation of ToxR production by CsrA. A strain containing mutations in G3 only was created and tested for its response to NRES. The G3 mutant produced less ToxR than wild type, but still exhibited NRES-dependent ToxR production (Fig 4C, top panel); however, the amount of ToxR produced in this mutant was not sufficient to fully induce *ompU* expression, nor did it fully repress *ompT* expression (Fig. 4C, bottom panel). This suggests that the G3 motif is important for the NRES and CsrA-mediated regulation of ToxR levels. Mutating G4 in addition to G3 further decreased NRES-dependent ToxR production (Fig. 4C, top panel), and no OmpU was detectable in response to NRES in this strain (Fig. 4C, lower panel). Mutating motifs G4 and G5 together significantly lowered the amount of ToxR produced overall, and the level of ToxR remained low, even when NRES was added to the growth medium (Fig. 4C, top panel). As expected, no switch from OmpT to OmpU was observed in this mutant in response to NRES (Fig. 4C, lower panel). Thus, the GGA motifs play central roles in regulation of ToxR synthesis by CsrA. CsrA dimers may bridge sequences spaced some distance apart within the RNA by binding two motifs at a time.

**Figure 4.**
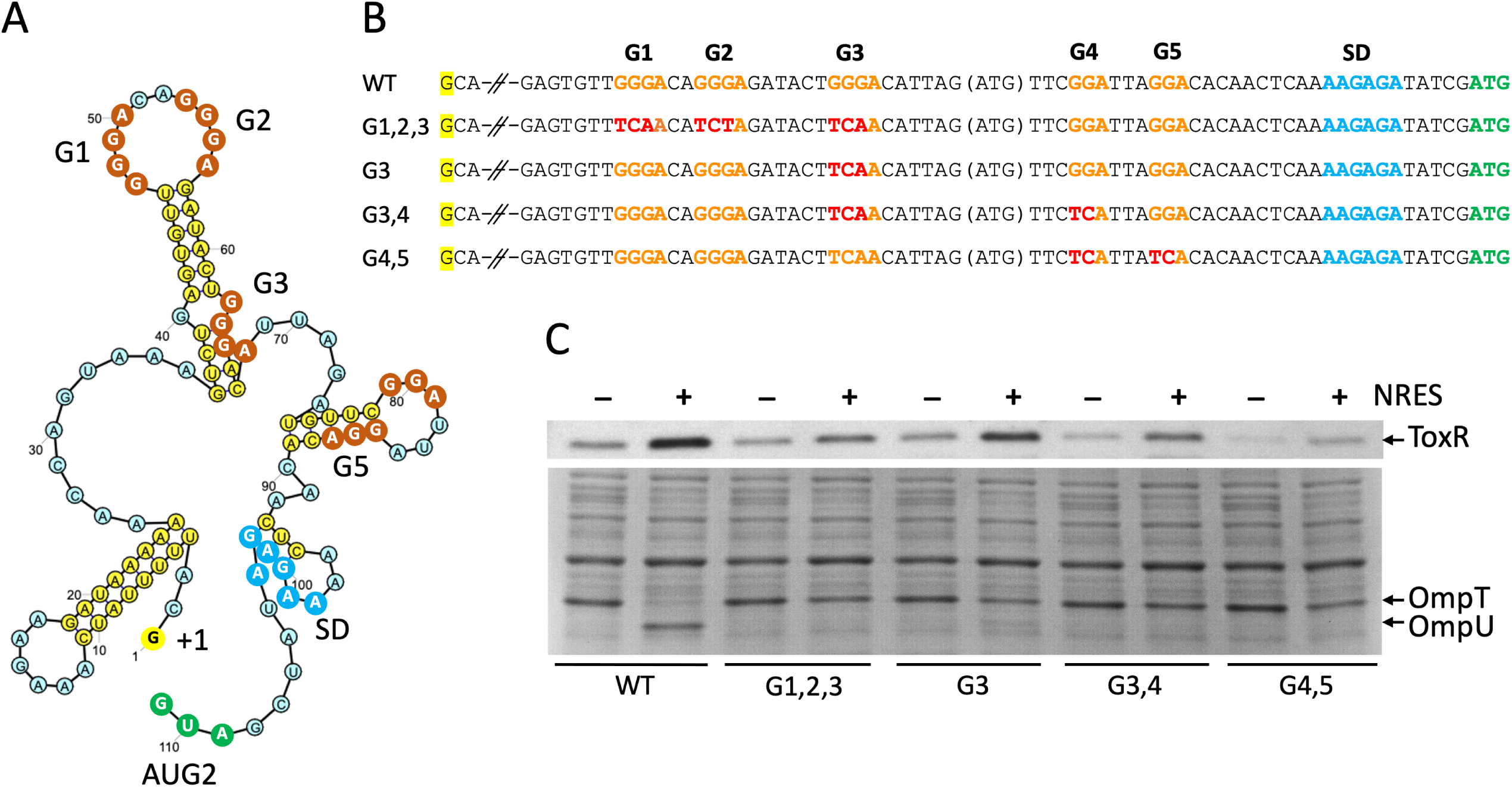
(G)GGA motifs in the *toxR* 5’UTR are essential for regulation of ToxR synthesis by CsrA and NRES. (A) Predicted structure of the 5’UTR RNA sequence using mfold to fold the 111 nucleotide segment (59) and Ribosketch to illustrate the secondary structure features. The 5’ nucleotide (+1) in the transcript sequence is highlighted in yellow, the (G)GGA motifs, numbered G1-G5 in the 5’-3’ direction, are shown with orange circles, the SD sequence is indicated with dark blue circles, and the AUG translational start is indicated with green circles. (B) Alignment of the 5’ UTR region containing the (G)GGA motifs (orange) from each of the mutants tested showing the specific mutations (red) of each strain compared with the wild-type sequence. The 5’ nucleotide (+1) in the transcript sequence is highlighted in yellow, and the caesura represents intervening sequence not depicted in the figure. (C) Strains N16961 (WT), N*toxR*.G1,2,3 (G1,2,3), N*toxR*.G3 (G3), N*toxR*.G3,4 (G3,4), and N*toxR*.G4,5 (G4,5) were grown to mid-logarithmic phase in T medium with or without 12.5 mM NRES mix. Cells were harvested, and whole cell preparations were resolved by SDS-PAGE and immunoblotted using polyclonal anti-ToxR antisera (top panel) or stained with Coomassie Blue to visualize the Omp proteins (bottom panel).

To test whether the low level of ToxR protein produced in some of the GGA motif mutants was functional, activation by bile was assessed. With the exception of the G4,5 mutant, the basal amount of ToxR was activated by cholate to produce a wild-type level of OmpU (Fig. S4D). A less robust response to cholate was observed in the G4,5 mutant, which also had the lowest basal expression of ToxR. It is possible that, being so close to the SD, the G4,5 mutations interfered with some aspect of translation.

### CsrA increases the level of *toxR* mRNA in NRES by stabilizing the transcript

We showed previously that the addition of NRES to cultures of *V. cholerae* growing in minimal medium caused the level of the *toxR* transcript to increase (26). To determine whether CsrA is required for the NRES-mediated increase in the *toxR* transcript, the strains were grown in minimal medium with or without NRES, and *toxR* mRNA levels were determined using RT-qPCR assays. The level of the *toxR* transcript increased almost 4-fold in the wild type in the presence of NRES (Fig. 5A), whereas no statistically significant increase in *toxR* mRNA was observed in the *csrA* mutant (Fig. 5A), demonstrating that CsrA is required for the increase in the *toxR* transcript level in response to NRES.

**Figure 5.**
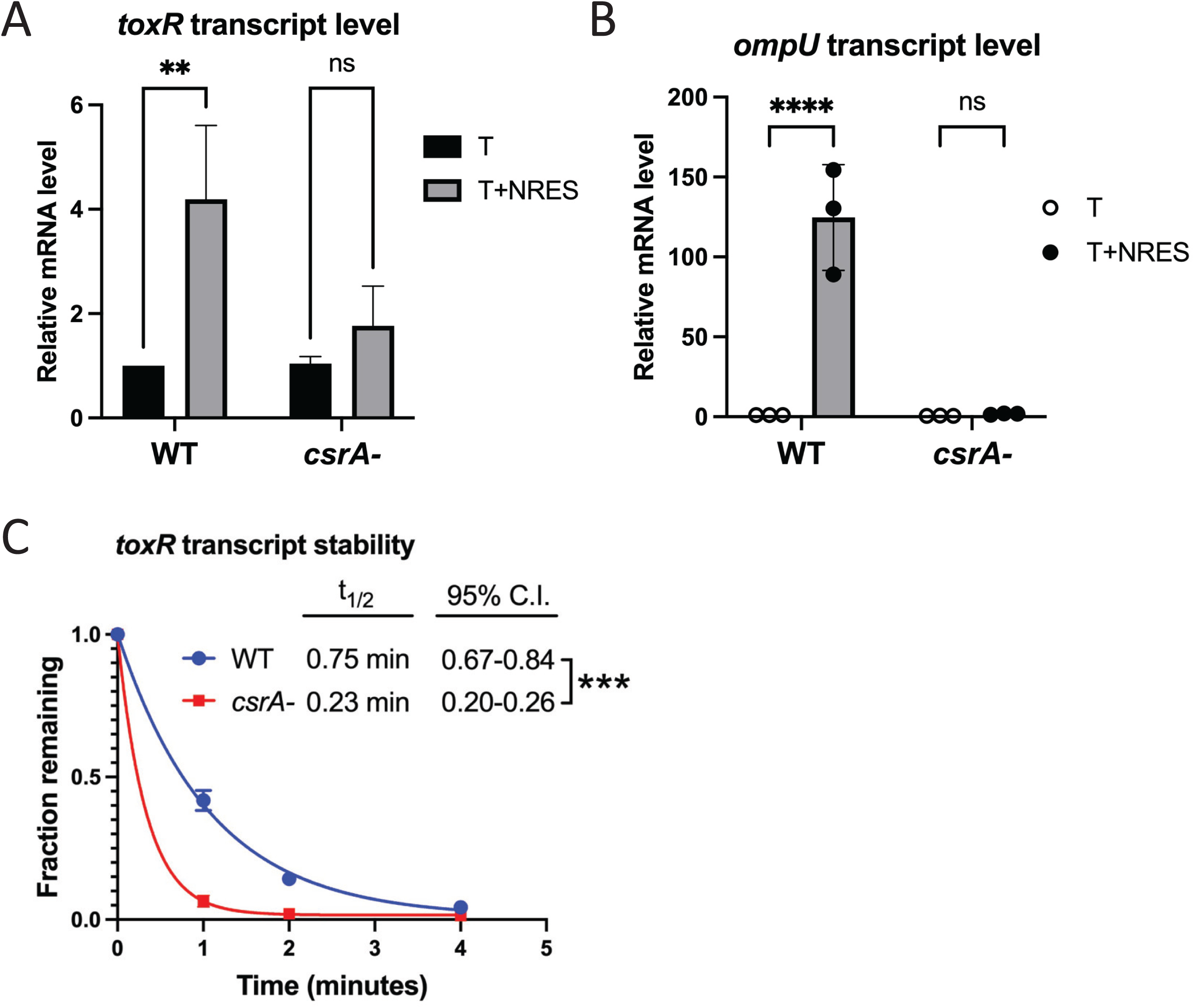
Both the amount of *toxR* mRNA in NRES and the stability of the *toxR* transcript are increased by CsrA. (A) Strains N16961 (WT) and N*csrA*.R6H (*csrA-*) were grown overnight in LB medium and diluted 1:100 into T medium with or without 12.5 mM NRES mix. RNA was isolated from cultures grown to mid-logarithmic phase, and the *toxR* (A) mRNA level was assessed by RT-qPCR with normalization to an internal control, *atpI*. Expression levels are shown relative to the level in the wild-type strain without NRES. The black bars represent expression in the absence of NRES, whereas the gray bars represent expression in the presence of NRES. The *P* values for significance were determined from three independent experiments, as described in the Methods section (**, *P* < 0.01; ns, not statistically significant). (B) The *ompU* mRNA level was determined as described above using the same RNA preparations as in (A). Expression levels are shown relative to the level in the wild-type strain without NRES. The open circles represent expression in the absence of NRES, whereas the closed circles represent expression in the presence of NRES. The gray bar represents the median relative mRNA level of the data points plotted. The *P* values for significance were determined from three independent experiments, as described in the Methods section (****, *P*<0.0001; ns, not significant). (C) Strains N16961 (WT) and N*csrA*.R6H (*csrA-*) were grown overnight in LB medium and diluted 1:100 into T medium with or without 12.5 mM NRES mix. Cultures were grown to mid-logarithmic phase, and rifampicin was added to terminate transcription. Samples were collected immediately (t=0) and at 1, 2, and 4 minutes post-rifampicin addition, and the amount of the *toxR* transcript at each time point was measured using RT-qPCR. The fraction remaining at each time point is shown relative to the level at t=0 in each strain. The half-life (t_1/2_) and 95% confidence interval (95% C.I.) were determined using nonlinear regression, one-phase decay analysis, and significance was determined by the extra sum-of-squares F test (***, *P* < 0.001).

ToxR is an activator of *ompU* transcription, and the increase in *toxR* mRNA in response to NRES correlated with a very large increase in the level of *ompU* mRNA (26). To verify that CsrA is also required for the transcriptional activation of *ompU* in response to NRES, the level of *ompU* was evaluated in the *csrA* mutant grown in NRES. In contrast to the wild-type strain, there was no statistically significant increase in the *ompU* transcript level in response to NRES in the *csrA* mutant, demonstrating an absolute requirement for CsrA in the NRES-induced *ompU* transcription activation by ToxR (Fig. 5B). Thus, CsrA is critical for increasing both the *toxR* and *ompU* transcript levels in response to NRES.

Since CsrA can bind to the *toxR* transcript directly, it is likely that CsrA increases *toxR* mRNA levels by binding to the *toxR* 5’UTR and protecting the transcript from degradation. To test whether CsrA plays a role in stabilizing the *toxR* transcript, the stability of the *toxR* transcript was assessed by measuring the decline in *toxR* mRNA in the wild-type and *csrA* mutant strains following addition of rifampicin, which prevents new synthesis of *toxR* transcripts (Fig. 5C). With a half-life of less than a minute (half-life = 0.75 min) in the wild-type strain, the *toxR* transcript appears to be relatively unstable, even in the presence of CsrA; however, in the *csrA* mutant, the half-life of the *toxR* transcript was further decreased by more than 3-fold (half-life = 0.23 min)(Fig. 5C), indicating that CsrA plays an important role in stabilizing *toxR* mRNA.

### CsrA stimulates *toxR* translation in a cell-free, in vitro protein synthesis assay

To determine whether CsrA also affects translation of the ToxR protein, we carried out a cell-free protein synthesis assay of ToxR in the presence and absence of CsrA. The template encompassed the *toxR* transcriptional start site (+1) and 5’UTR, together with the native SD and entire *toxR* open reading frame. To eliminate any regulatory effects of the *toxR* promoter in the assay, expression of *toxR* was driven from a T7 promoter, and no native *toxR* promoter sequences were included (Fig. 6A). An RNase inhibitor was added to the cell-free protein synthesis reaction to exclude any effects of CsrA on mRNA stability in the in vitro translation assay. Addition of purified CsrA to the in vitro transcription-translation assay greatly enhanced ToxR production (Fig. 6B, upper panel), consistent with CsrA having a direct effect on translation of the *toxR* transcript. In the absence of CsrA, there was no difference in the amount of ToxR produced with or without added glycerol (Fig. 6B), ruling out any stimulatory effects of glycerol in the CsrA protein storage buffer. No effects of CsrA on the synthesis of the control protein DHFR were detected, showing that CsrA does not generally promote T7 transcription or translation in the cell-free system (Fig. 6B, lower panel). Taken together, these results demonstrate that CsrA enhances translation of ToxR from a transcript containing only the *toxR* 5’UTR and open reading frame. Thus, the effects of CsrA on ToxR production appear to be two-fold: stabilization of the *toxR* mRNA and activation of *toxR* translation.

**Figure 6.**
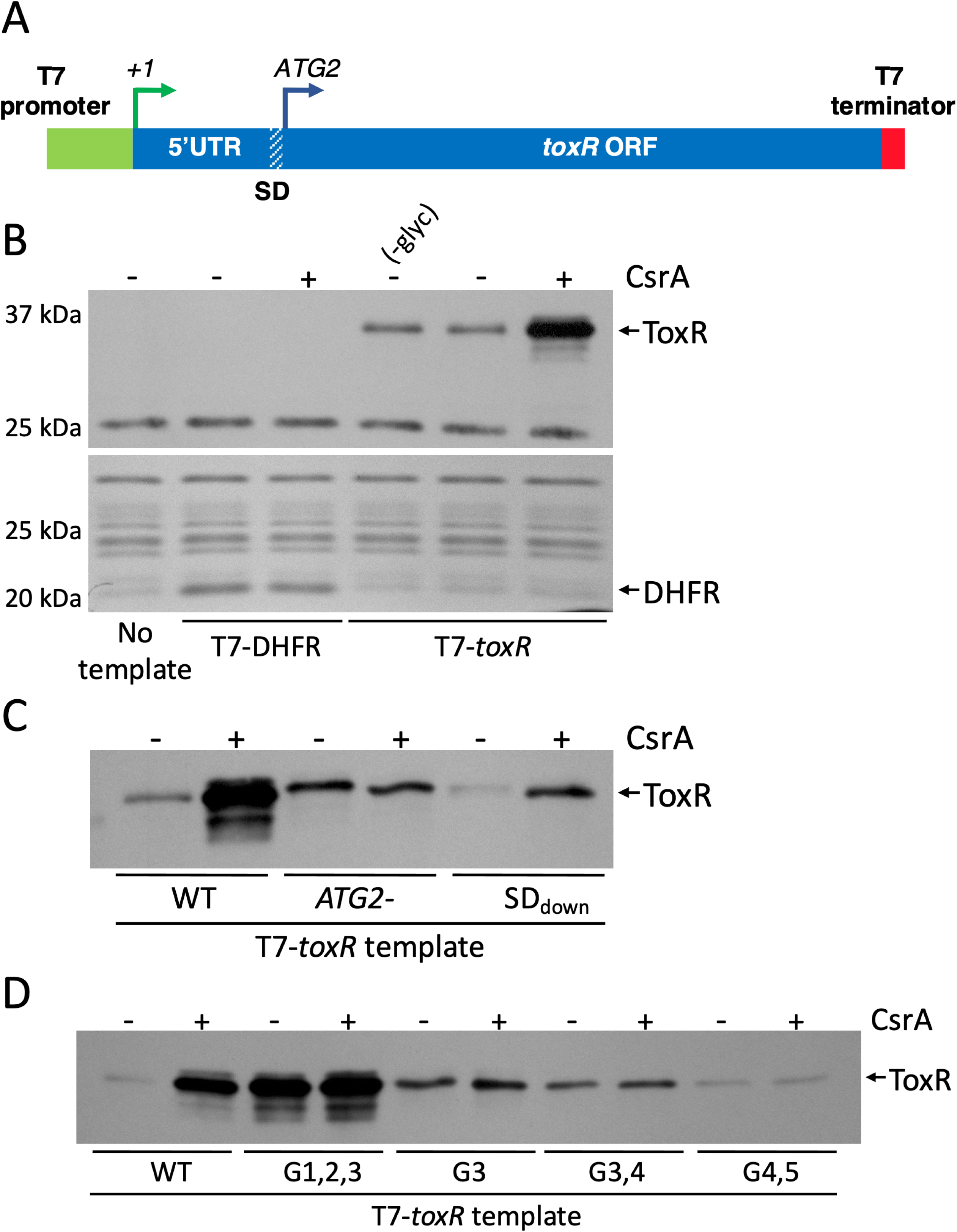
CsrA stimulates *toxR* translation in a cell-free, in vitro protein synthesis assay. (A) Diagram of the DNA template used for the cell-free in vitro transcription-translation assays. The template comprises the T7 polymerase promoter consensus sequence fused to the 5’UTR, SD, and coding sequences of the *toxR* gene, with a T7 transcriptional terminator at the 3’ end. (B) PURExpress cell-free, in vitro protein synthesis assays were carried out with or without purified CsrA. The relevant templates are listed below the images. The PURExpress reactions were resolved by SDS-PAGE and immunoblotted using polyclonal anti-ToxR antisera to visualize synthesis of ToxR (top panel) or stained with Coomassie Blue to visualize synthesis of the control protein DHFR (bottom panel). The relevant proteins are indicated with arrows on the right. (C) T7-*toxR* templates carrying *toxR* 5’UTR mutations (T7-*toxR*.M13L and T7-*toxR*.SDdown) shown to interfere with translation from the second in-frame ATG in vivo (Figs. 1 and 2) were compared with the wild-type T7-*toxR* template in the in vitro transcription-translation assay with and without purified CsrA. The PURExpress reactions were resolved by SDS-PAGE and immunoblotted using polyclonal anti-ToxR antisera to visualize synthesis of ToxR. The ToxR protein is indicated with an arrow on the right. (D) T7-*toxR* templates carrying mutations in the *toxR* 5’UTR GGA motifs were compared with the wild-type T7-*toxR* template in the in vitro transcription-translation assay with and without purified CsrA. The PURExpress reactions were resolved by SDS-PAGE and immunoblotted using polyclonal anti-ToxR antisera to visualize synthesis of ToxR. The ToxR protein is indicated with an arrow on the right.

The in vivo experiments in this study indicated that ToxR is translated from the second in-frame ATG. To test whether this is observed also in the in vitro system, an in vitro transcript was generated with a mutation in ATG2 to test whether any ToxR protein is translated from the first in-frame ATG. As shown in Figure 6C, a ToxR protein was made, but the synthesis of ToxR from the first ATG was not responsive to CsrA, similar to the in vivo result, showing no effects of NRES on the production of ATG1-derived ToxR (Fig. S1). The SD sequence identified as being essential for in vivo translation of *toxR* (Fig. 1D) was shown to be critical also for in vitro translation: the level of ToxR translated from the SDdown mutant transcript was much lower than from the wild-type transcript (Fig. 6C); however, as observed in vivo (Fig. 1D), translation was responsive to CsrA, as a significant increase in ToxR levels was detected when CsrA was added to the reaction (Fig. 6C). Thus, the SDdown mutations did not interfere with CsrA regulation per se, but with translation efficiency generally, as would be expected from introducing deleterious mutations into the SD sequence.

To test the importance of the *toxR* 5’UTR GGA motifs for efficiency of translation in vitro, templates were generated with mutations in these motifs. Mutating the G3 motif, either alone or in combination with the G4 motif, was sufficient to abolish the CsrA-mediated increase in ToxR synthesis (Fig. 6D). In both cases, the mutant transcripts produced above base-line levels of ToxR, but in equal measure with and without CsrA (Fig. 6D), showing that translation is independent of CsrA. The baseline ToxR production from the G4,5 mutant template was very low, as expected based on the in vivo results, and no effects of adding CsrA to the in vitro synthesis assay were detected. Interestingly, synthesis of ToxR from the G1,2,3 mutant transcript showed an unexpected phenotype in the cell-free assay. Although *toxR* translation did not appear to be greatly stimulated by CsrA, the basal level of ToxR produced both with and without CsrA was very high. This suggests that the mutations may have eliminated an inhibitory element in the *toxR* mRNA in vitro.

## DISCUSSION

CsrA is a major regulator of virulence gene expression in *V. cholerae* and is essential for pathogenesis in a mammalian intestinal infection model system (34). We have identified several targets of *V. cholerae* CsrA regulation with documented roles in virulence, including the transcriptional regulators VarA (45, 60), AphA (35, 61), and ToxR (34). Interestingly, the mechanism of regulation in all three cases appears to be direct and positive, with CsrA binding directly to the target transcript and causing protein synthesis to increase. This is not a common mechanism among the characterized modes of regulation by CsrA, and therefore there is a lack of clearly defined paradigms for comparison (62). In this study, we demonstrate that CsrA increases ToxR levels in the cell by increasing the *toxR* mRNA stability and abundance, and by increasing the efficiency of ToxR translation.

CsrA typically regulates its targets by binding to the 5’UTR of the mRNA, but the *toxR* 5’UTR had not been clearly defined. Here, we delineate the 5’UTR of *toxR* by establishing the second in-frame ATG as the main translation start. While the amino acid sequence encoded between the first and second in-frame ATGs was not critical for ToxR function, the nucleotide sequence houses important regulatory motifs, including two GGA motifs with roles in CsrA-mediated regulation, and a SD sequence with the appropriate spacing (4-9 nucleotides between SD and the adenine of the ATG start codon) (63) upstream of ATG2. This SD sequence is a relatively poor match to the consensus SD sequence in *E. coli* (64) and does not contain the central GGA sequence often present in typical SD sequences. Given that CsrA is a positive regulator of ToxR synthesis, this is not surprising; SD sequences that contain GGA may represent a target for CsrA binding, which would block, rather than increase, translation (46, 65). In fact, when we altered the identified SD to match the consensus sequence, *toxR* translation decreased, possibly because a new CsrA binding site was introduced.

Although ToxR was primarily translated from the second in-frame ATG, a small amount of ToxR was produced from the first in-frame ATG. It is not yet apparent whether this longer form of ToxR has any relevance for *V. cholerae* virulence gene regulation. Alignments of *V. cholerae* ToxR with ToxR proteins from other *Vibrio* species show that other ToxR homologs generally do not encode these extra N-terminal amino acids, but rather initiate at the codon corresponding to ATG2 (53). The longer variant of ToxR was poorly activated by bile, resulting in a very low level of OmpU production. Since bile is a potent activator of ToxR activity, promoting robust *ompU* expression even when ToxR levels are extremely low, the low OmpU level suggests that the addition of extra amino acids to the N-terminus of ToxR is deleterious to its function. Nevertheless, conditions may exist under which this variant plays a role in the regulation or activity of ToxR.

The CsrA-mediated increase in ToxR protein levels in response to NRES correlated with an increase in the amount of *toxR* mRNA in the cell. Further, the stability of *toxR* mRNA was greatly increased by CsrA, likely the result of a direct interaction between CsrA and the *toxR* transcript. Binding of CsrA may stabilize the *toxR* mRNA by shielding it from nucleases, either by having a binding site that overlaps with a nuclease cleavage site, as was shown for the sRNA spot 42 in *E. coli* (66), or by stabilizing a secondary structure of the transcript that reduces nuclease access to the transcript, as exemplified by the protection of the *E. coli flhDC* transcript through CsrA binding at the 5’ end of the transcript (42). It is also possible that the *toxR* transcript is more stable due to an increase in translation in the presence of CsrA. Increased ribosomal occupancy may occlude nuclease cleavage sites, and higher translation rates appear to have a protective effect on mRNA stability (67, 68).

In *Salmonella enterica* serovar Typhimurium, CsrA appears to act as a stabilizer of RNA transcripts globally, with the mean half-life of transcripts nearly doubling in the presence of CsrA. Of note, however, the increase in overall stability was not necessarily associated with an increase in overall abundance of mRNA in the cell, and the effects of CsrA were still largely negative on gene expression (39). This could be due to other factors besides mRNA stability that dictate transcript abundance, including rate of transcription and rate of dilution of transcripts during cell division. A similar CsrA-dependent increase in overall mRNA stability was observed in *E. coli* when comparing global transcript stability in the presence or absence of CsrA (69). As in the *Salmonella* study, the increased transcript stability in the presence of CsrA did not necessarily correlate positively with mRNA abundance; however, a significant number of CsrA-stabilized transcripts also showed an increase in the total mRNA level, suggesting that CsrA-mediated stabilization of mRNAs may be relevant for specific transcripts and could be a more common mechanism of regulation that previously understood (69). We observed a three-fold increase in the *toxR* mRNA half-life, and a four-fold increase in transcript abundance in the presence of CsrA, consistent with a role for CsrA in increasing transcript abundance by stabilizing the RNA.

The role of the characteristic high-affinity CsrA binding site, consisting of a GGA sequence exposed in the loop of a hairpin (stem-loop) structure in either sRNAs or in the 5’UTR of mRNA targets, is well documented in several species, including *E. coli* and *Pseudomonas* (54, 70–73). In this study, we provide direct evidence that GGA motifs are involved in the interaction of CsrA with RNA targets also in *V. cholerae*. Oligonucleotides representing a portion of the *V. cholerae* CsrB sRNA were used to demonstrate that optimal CsrA binding requires two GGA motifs located in the loops of adjacent stem-loop structures within the RNA. Mutation of either GGA sequence eliminated CsrA binding, even though the predicted stem-loop structures themselves remained intact. This was surprising, given that most previous studies analyzing the in vitro interactions between CsrA (or CsrA homologs, such as RsmE) and substrate oligonucleotides have shown that CsrA can bind to single RNA and DNA hairpin oligonucleotides with only one GGA motif. In NMR solution structures of *Pseudomonas fluorescens* RsmE complexed with an RNA substrate, RsmE bound to the RNA oligonucleotide in a 1:1 molar ratio, resulting in binding of two independent RNA molecules per RsmE dimer (71). In *Bacillus subtilis,* the CsrA dimer bound two DNA hairpin (but not linear) oligonucleotides, each with a single GGA motif, but the oligonucleotides did not need to be connected for binding to occur (74). Similarly, Mercante et al, 2009 (75) showed in REMSA experiments that the *E. coli* CsrA dimer contains two independent RNA-binding surfaces and was able bind to two separate RNA oligonucleotides simultaneously; however, they also showed that the CsrA dimer could bridge two high-affinity binding sites within a single RNA oligonucleotide, although CsrA was still able to bind if one of the GGA motifs was mutated. This is in contrast to our findings in *V. cholerae*, which showed that both GGA motifs are required for binding. It is not clear whether these discrepancies reflect differences in the assay conditions or real differences in the binding requirements for *V. cholerae* CsrA. Nevertheless, it is common for both *E. coli* and *V. cholerae* CsrA targets to have more than one GGA motif in the 5’UTR of the transcript (this study; (45, 72, 75)), and the ability of *E. coli* CsrA to bridge two sites has been shown to be required for efficient post-transcriptional regulation (72, 75). Thus, the binding of the CsrA dimer to two sites within the same RNA strand may be a key regulatory feature of CsrA proteins in general. Importantly, the presence of multiple GGA motifs in the *toxR* 5’UTR, together with the observed requirement for two binding motifs in a single RNA strand for efficient binding of *V. cholerae* CsrA to an RNA substrate, suggests that each bound *V. cholerae* CsrA dimer probably binds two sites at a time. Depending on which sites are occupied, and whether the bridged sites are adjacent or spaced apart, this could have substantial effects on the secondary structure of the *toxR* 5’UTR, and thus on the accessibility of the ribosome to the SD sequence and the initiator AUG.

The functional importance of the *toxR* 5’UTR GGA motifs was demonstrated using mutational analysis, both in vivo and in vitro. Mutating the GGA motifs in vivo, particularly when two or more motifs were mutated at the same time, was associated with significant defects in the NRES response. In most of the mutants tested, the amount of ToxR produced failed to increase to the level required to induce Omp switching in NRES. As expected, the cholate response was largely unaffected by the GGA mutations, since the cholate response does not require CsrA. Only one mutant, G4,5, showed less efficient Omp switching in response to cholate, possibly due to the proximity of the mutations to the start of translation, where they might interfere with ribosome loading. In support of this, the level of ToxR in this strain was found to be significantly lower in the G4,5 mutant than in the wild-type strain, regardless of the growth condition. A similar defect in the amount of ToxR produced from a template containing the G4,5 mutations compared with the wild-type template was observed in vitro, supporting the hypothesis that these mutations had a general deleterious effect on translation.

The in vitro assays showed clearly that CsrA stimulates translation from the wild-type *toxR* template: the amount of ToxR produced in the cell-free reaction was greatly increased by the addition of CsrA. We hypothesize that, in the absence of CsrA, the 5’UTR folds into an inhibitory structure that prevents ribosome loading. CsrA binding may cause the 5’UTR to adopt a secondary structure that releases the ribosome binding sequence. Evidence for this comes from results showing that the requirement for CsrA in an in vitro translation assay is circumvented by mutating the first three GGA motifs in the 5’UTR template. This suggests that, at least in vitro, the mutated template may be unable to adopt the inhibitory structure, thus rendering the template much more favorable for translation initiation, even in the absence of CsrA. Interestingly, this was not the case in vivo, where these same mutations abolished, rather than increased, production of ToxR in response to NRES; however, in vivo, there may be other factors that come into play as well, including the effects of CsrA on the stability of the transcript. While the G1,2,3 mutations may prevent formation of an inhibitory secondary structure, the resulting loss of CsrA binding may render the transcript unstable in vivo. Transcript instability would not be relevant in vitro, as there are no RNases present, and the template is present in excess.

*V. cholerae* is an aquatic organism with remarkable adaptability, and its success as a pathogen stems in large part from its ability to negotiate substantial environmental shifts by adjusting its gene expression profile. The host intestinal environment poses a unique set of challenges, as well as opportunities, for *V. cholerae*. The lumen of the small intestine contains a number of toxic compounds that are harmful to the invading bacteria, and a rapid response to these compounds is essential for survival. The immediate activation of ToxR activity upon encountering intestinal bile allows *V. cholerae* to quickly make changes in the composition of its outer membrane to ensure that bile is excluded from the cell. The intestinal environment is also nutrient-rich, and coordinating factors required for colonization with the presence of nutrients ensures a successful infection. Increasing the synthesis of the ToxR protein in response to carbon sources such as amino acids and intestinal mucin (26) represents a longer-term shift in bacterial physiology to maximize infection potential in a favorable environment. In this study we demonstrate that CsrA, by binding directly to the *toxR* transcript and increasing both the stability of the transcript and translation of the ToxR protein, is a master regulator of an environmental response that ultimately results in colonization and pathogenesis by this important human pathogen.

## MATERIALS AND METHODS

### Bacterial strains and plasmids, and growth conditions

Bacterial strains and plasmids used in this study are listed in Table S1. All strains were maintained at –80°C in tryptic soy broth (TSB) plus 20% glycerol. Strains were routinely grown at 37°C in either Luria-Bertani (LB) broth (w/vol: 1% tryptone, 0.5% yeast extract, 1% NaCl) (76) or in T medium (77) modified to contain 0.2% w/vol sucrose, 20 μM FeSO_4_, and a mixture of vitamins (recipe for 100X VA vitamin solution at http://www.genome.wisc.edu/resources/protocols/ezmedium.htm). Amino acids (L-asparagine, L-arginine, L-glutamic acid, L-serine)(Sigma-Aldrich, St. Louis, MO) for the NRES supplement were dissolved in water and used at a final concentration of 12.5 mM total amino acids (3.125 mM each amino acid). Sodium cholate hydrate (Sigma-Aldrich) was used at 0.1% (w/vol). Antibiotics were used at the following concentrations for *E. coli* strains: 250 μg of carbenicillin per ml, 50 μg of kanamycin per ml, and 30 μg of chloramphenicol per ml. For *V. cholerae* strains, the concentrations used were 125 μg of carbenicillin per ml, 25 μg of kanamycin per ml, 6 μg of chloramphenicol per ml, 30 μg of gentamicin per ml, and 20 μg polymyxin B per ml.

Plasmid clones were transferred to *V. cholerae* strains by electroporation or bacterial conjugation, as described previously (78)

### Polymerase chain reaction (PCR)

The oligonucleotide primers for PCR were purchased from Sigma-Aldrich and from Invitrogen (Carlsbad, CA). Primers used for cloning are listed in Table S2. PCR was performed using KOD Hot Start DNA polymerase (Novagen®, MilliporeSigma, Burlington, MA) to create fragments for cloning, and *Taq* DNA polymerase (Qiagen, Valencia, CA) for all general verification purposes. PCR reactions were carried out according to the manufacturer’s instructions, using the primers described below. Unless otherwise indicated, the template for PCR was bacterial whole cell suspensions in water, boiled for 5 minutes. All clones derived from PCR products were verified by DNA sequencing.

### Sequence Analysis

DNA sequencing was performed by the University of Texas Institute for Cellular and Molecular Biology DNA Core Facility using the capillary-based 3730 DNA Analyzer from Applied Biosystems (Thermo Fisher Scientific, Waltham, MA). Analysis of DNA sequences was carried out using MacVector 15.1.5. BLAST searches and other bioinformatics analyses were done using the National Center for Biotechnology Information (NCBI) and the Comprehensive Microbial Resource (CMR) databases. Pairwise alignments were carried out using ClustalW from within MacVector 18.6.4. RNA structure modeling was performed using the mfold nucleic acid folding program (59) through the UNAFold Web Server (http://www.unafold.org/RNA_form.php). Identification and analysis of the 8 *V. cholerae* 16s rRNA gene sequences was carried out using a Microbial BLASTN search (https://blast.ncbi.nlm.nih.gov/Blast.cgi) of the *Vibrio cholerae* O1 biovar El Tor str. N16961 (taxid:243277) genome using the *E. coli rrsA* gene sequence as the query.

### Construction of *V. cholerae* chromosomal mutations

To create chromosomal mutations in the 5’UTR of *toxR*, two fragments with a short overlap were generated by PCR using the primers listed in Table S2. The first fragment was amplified using primer htpG1 together with the reverse mutagenic primer, while the second fragment was amplified using the forward mutagenic primer together with primer toxS2. The overlapping fragments were then used as the template to generate a splice overlap extension (SOE) PCR product using primers htpG1 and toxS2. The final PCR fragment containing the mutation(s) was cloned into the SmaI site of pCVD442N to yield the allelic exchange constructs listed in Table S1 (pAMS33-pAMS44). To introduce a V5 tag at the carboxy terminus of ToxS, overlapping fragments containing the V5 coding sequences and 3 glycine linker codons fused in frame to the 3’ end of *toxS* were generated using primer pairs toxS-V5.1/ toxS-V5.2 and toxS-V5.3/ toxS-V5.4. The overlapping fragments were used as the template for SOE PCR using primer pair toxS-V5.1/ toxS-V5.4 to generate the toxS-V5 allele. The mutagenic fragment was cloned into the SmaI site of pCVD442N to yield the allelic exchange construct pStoxS-V5.

Allelic exchange constructs were transferred to *V. cholerae* strains via bacterial conjugation, and allelic exchange was carried out as described previously (79, 80).

### SDS-polyacrylamide gel electrophoresis (PAGE) and immunoblotting

Cultures were grown overnight in LB medium and diluted 1:100 into T medium, supplemented as described above and in the figure legends. The diluted were grown to mid-log phase (OD_650_ ≈ 0.4-0.8), and samples containing an equivalent number of cells were resuspended in Laemmli solubilization buffer (81). Whole cell extracts were resolved by SDS-10% PAGE and visualized by Coomassie Brilliant Blue staining or electroblotted for 1.5 hr at 45V onto Hybond^TM^ ECL^TM^ nitrocellulose (Amersham Pharmacia Biotech, Little Chalfont, Buckinghamshire, England). The positions of OmpT and OmpU were determined through comparisons with previously published Coomassie-stained gel analyses of *ompT* and *ompU* mutants (82), and by OmpT and OmpU immunodetection assays using anti-OmpT antisera (diluted 1:5000)(generous gift of J. W. Peterson), which reacts with both OmpT and OmpU (82), and HRP-conjugated goat anti-rabbit IgG (Bio-Rad Laboratories, Hercules, CA) secondary antibody (diluted 1:5000).

Immunodetection of ToxR was carried out using rabbit polyclonal anti-ToxR antiserum (diluted 1:4000) (generous gift of R. K. Taylor), and HRP-conjugated goat anti-rabbit IgG secondary antibody (diluted 1:5000). The V5 epitope tag was detected using mouse monoclonal anti-V5 antibodies (Sigma-Aldrich) and HRP-conjugated goat anti-mouse IgG (Bio-Rad laboratories) secondary antibody (diluted 1:5000). Equal loading of the samples for immunoblotting was confirmed by evaluating the corresponding Coomassie-stained gel. Each gel or blot shown in the Results section is representative of at least three independent experiments.

### Protein Expression and Purification

The Tobacco Etch Virus (TEV) protease was purified as follows: The TEV expression strain [the Codon plus DE3 pRIL strain (Agilent Technologies, Santa Clara, CA) harboring plasmid pRK793 (83) was grown at 37 °C overnight in ZYP-0.8G medium (ZYP medium (84) supplemented with 0.8% glucose) containing 100 μg/ml of carbenicillin and 50 μg/ml of chloramphenicol. The overnight culture was diluted 1:100 into 25 ml of TB with 200 μg/ml of carbenicillin and incubated at 37 °C until growth reached an OD_600_ of ∼4. This culture was then used to inoculate the 250 ml production culture (1:10) of TB supplemented with 2 mM MgSO4 and 50 μg/ml carbenicillin. The culture was incubated at 37 °C until growth reached an OD_600_ of 0.8, at which point 200 μM IPTG and 5% glycerol were added. Following overnight incubation at 18 °C, cells were centrifuged at 4000 xg for 25 minutes and resuspended in 20 ml of lysis buffer (50 mM Tris pH 7.2, 150 mM NaCl, 20 mM imidazole, with one Complete EDTA free protease inhibitor tablet (Roche, Basel, Switzerland)). The cells were lysed with 3 passes through a French press, and the lysate was centrifuged at 100,000 xg for 45 minutes. The supernatant was removed and filtered using a 0.45 μm filter. The protein was captured on a HisTrap™column (GE Healthcare, Chicago, IL) equilibrated with wash buffer (50 mM Tris pH 8.5, 20 mM imidazole, 500 mM NaCl). The column was washed with 30 CV of wash buffer, then with 9 CV of 9% elution buffer (50 mM Tris pH 8.0, 500 mM imidazole, 500 mM NaCl). The elution was performed with a gradient to 100% elution buffer over 12 CV. The relevant fractions were pooled and dialyzed against 4 L of dialysis buffer (25 mM Tris pH 7.8, 150 mM NaCl, 1 mM EDTA, 2 mM DTT, 10% glycerol) overnight at 4 °C. After addition of 5% glycerol to the protein, the protein was aliquoted, flash frozen, and stored at -80 °C.

To purify *V. cholerae* CsrA, a codon-optimized *V. cholerae csrA* open reading frame encoding additionally an N-terminal His tag and containing a TEV cleavage site was synthesized by IDT (Coralville, IA). The construct includes NcoI and BamHI restriction sites for cloning and the M13 forward (-21) and reverse primer sequences for PCR. The *csrA* construct was PCR amplified, digested with NcoI/BamHI, and ligated into NcoI/BamHI-digested, calf intestinal phosphatase-treated pET16b vector (Novagen®, MilliporeSigma) using Quick Ligase (New England Biolabs [NEB], Ipswich, MA). The resulting plasmid, pHTCsrA, was verified by sequencing and transformed into BL21 DE3 cells for expression. For expression, a starter culture, inoculated from a frozen stock, was started in 2 ml of ZYP-0.8G medium grown overnight at 30°C. After growth overnight, the starter culture was diluted 1:100 into a small culture (25 mL) of TB and incubated at 37°C until the culture reached an OD_600_ of about 5. The culture was then diluted 1:10 into a 250 mL production culture that was incubated at 37°C until the culture reached an OD_600_ between 2 and 3. The culture was induced with 200 μM IPTG and 5% glycerol was added. Following incubation of the culture overnight at 25°C, cells were collected by centrifugation at 4500 xg at 4°C. The cells were resuspended in 20 mL lysis buffer composed of wash buffer (50 mM KPO_4_ pH 8.0, 10 mM imidazole, 300 mM NaCl) with a Complete EDTA free Protease Inhibitor Tablet (Roche). The cells were lysed by three passes through a French pressure cell. The lysate was clarified at 100,000 xg for 45 minutes. The clarified lysate was filtered with a 0.45 μm filter and the protein was captured using a HisTrap™ column. The column was washed with 10 column volumes (CV) of wash buffer, 10 CV of high salt wash buffer (50 mM KPO_4_ pH 8, 1 M NaCl, 20mM imidazole), then with 10 CV of wash buffer. The protein was eluted with a 0-99% gradient of elution buffer (50 mM KPO_4_ pH 8.0, 250 mM imidazole, 300 mM NaCl). The relevant fractions were pooled, incubated with TEV protease, and dialyzed overnight against cleavage buffer (50 mM KPO_4_ pH 8.0, 150 mM NaCl, 2 mM DTT). The protease, the cleaved His tag, and any uncleaved protein were removed by flowing the dialysate, supplemented with 80 mM imidazole, over a HisTrap™ column. The flow-through was collected, concentrated, and the final purification was performed with a S75 16/600 gel filtration column (GE Healthcare) with gel filtration buffer (50 mM KPO_4_, 50 mM NaCl). The relevant fractions were collected, and the protein was concentrated as required.

### RNA electrophoretic mobility shift assay (REMSA)

RNA oligonucleotides representing partial sequences of CsrB with or without mutations were designed and tested for the most favorable folding using mfold (59). The partial CsrB RNA oligonucleotides (Table S3) were synthesized with a 3’ biotinylation modification (Sigma-Aldrich) and used in REMSA assays with purified CsrA. The 111-base RNA oligonucleotide corresponding to the 5’UTR of *toxR* (Table S3, oligo *toxR* 5’UTR), starting with the first base of transcription and ending with the second in-frame AUG (Fig 4A), was synthesized as an Ultramer® RNA oligonucleotide with a 5’ biotinylation modification (IDT). The oligonucleotides were diluted to 10 nM in 1x TE buffer and heated to 85°C for 4 minutes, followed by cooling to room temperature and storage on ice prior to the binding reactions. The binding reactions were carried out using the LightShift™ Chemiluminescent RNA EMSA Kit (Thermo Fisher Scientific), as per the manufacturer’s instructions. The final CsrB oligonucleotide concentrations used in the binding reactions ranged from 0.01 nM to 1 nM, as indicated in the relevant figure legends, and the final *toxR* 5’UTR oligonucleotide concentration was 1 nM. The concentration of purified CsrA ranged from 0-640 nM, as indicated in the relevant figure legends. The binding reactions were incubated for 30 min at 37°C, and the reactions were then resolved on an RNase-free, non-denaturing 0.5X TBE, 10% polyacrylamide gel, pre-run for 1 hour. Following electrophoresis, the gel was blotted to a BrightStar™-Plus Positively Charged Nylon Membrane (Thermo Fisher Scientific) using a semi-dry electroblotter for 20 min at 15 V. The blot was UV-crosslinked at 150 mJ and then developed according to the LightShift™ Chemiluminescent RNA EMSA Kit instructions to visualize the position of the biotinylated RNA oligonucleotide. Note: the CsrB oligo gave a much brighter signal at 1 nM than the other oligonucleotides at the same concentration, possibly due to more efficient labeling of this particular oligonucleotide. Therefore, CsrB oligo was used at a lower concentration (0.1 nM) in most experiments.

### RNA isolation and reverse transcriptase quantitative PCR (RT-qPCR)

Cultures were grown overnight in LB medium and diluted 1:100 into T medium supplemented with 0.2% sucrose, 20 μM FeSO_4_, and a mix of vitamins, as described above. The cultures were then divided, and the NRES solution was added to one set of cultures at a final concentration of 12.5 mM total amino acids. Cultures were grown to mid-log phase (OD_650_ ≈ 0.5) and then treated with an RNase-free, ice-cold solution of 95% absolute ethanol/5% phenol, pH 4.5, used at 20% vol/vol to preserve the RNA, and the cultures were kept on ice until further processing. Approximately 10^9^ cells per sample were collected by centrifugation and resuspended in 100 μL lysozyme solution (1 mg/mL lysozyme in Tris-EDTA buffer, pH 8.0). RNA was isolated using either RNA-Bee (Tel-Test, Inc., Friendswood, TX) or RNA-STAT 60 (Tel-Test, Inc.), as per the manufacturer’s instructions for isolation of total RNA from bacterial cells. Following purification, each RNA sample was DNaseI-treated using the TURBO DNA-*free*™ Kit (Invitrogen by Thermo Fisher Scientific), as per the manufacturer’s protocol. The RNA samples were resuspended in RNase-free water and quantified using an ND-1000 spectrophotometer (NanoDrop Technologies, Wilmington, DE). For RT-qPCR, 2 μg RNA per sample was used to generate cDNA using the High-Capacity cDNA Reverse Transcription Kit, and qPCR was carried out using Power SYBR® Green Master Mix, as per the manufacturer’s instructions (Thermo Fisher Scientific). The qPCR reactions were performed in an Applied Biosystems ViiA™ 7 Thermocycler using the following parameters: 50°C for 2 min, then 95°C for 10 min (hold stages), followed by 40 cycles of 95°C for 15 sec and 60°C for 1 min (PCR stages), with fluorescence recorded at the 60°C step. The relative expression levels were determined using the ΔΔCt method, where the Ct value is defined as the number of amplification cycles required to reach a fixed signal threshold. The reference gene used for normalization was *atpI*, which is not regulated by CsrA in *V. cholerae* (35). Two-way analysis of variance with Holm-Sídák corrections for multiple comparisons of gene expression in the wild-type strain N16961 (WT) versus the N*csrA*.R6H mutant (*csrA-*), in the presence or absence of NRES, was used to determine significance from the ΔCT values of three biological replicates (*, *P* < 0.05; **, *P* < 0.01; ***, *P* < 0.001), as described previously (85).

### RNA Stability Assay

Cultures were grown overnight in LB medium and diluted 1:100 into T medium supplemented with 0.2% sucrose, 20 μM FeSO_4_, and a mix of vitamins, as described above. The cultures were divided, the NRES solution (12.5 mM final amino acid concentration) was added to one set, and the cultures were grown to mid-log phase (OD_650_ ≈ 0.5). To stop transcription, 200 μg/mL rifampicin (Sigma-Aldrich) was added to the cultures and samples were collected immediately (t=0) and at 1, 2, and 4 minutes post-rifampicin addition. To preserve the RNA, samples were treated immediately upon collection with an RNase-free, ice-cold solution of 95% absolute ethanol/5% phenol, pH 4.5, used at 20% vol/vol and kept on ice until further processing. RNA isolation, cDNA synthesis, and RT-qPCR was caried out as described above, but using *rrsA* (16s rRNA) as the internal standard for approximation of overall cellular RNA stability for normalization purposes. The decay rate of the *toxR* transcript was determined by using the ΔΔCt method to calculate the fraction of *toxR* transcript remaining after each time point relative to t=0. Nonlinear regression, one-phase decay analysis in GraphPad Prism 10 was carried out to determine the half-life and confidence interval of the *toxR* transcript in each strain. The extra sum-of-squares F test was used to determine significance, as described previously (45).

### In vitro transcription-translation assay

In vitro, cell-free transcription-translation assays were carried out using the PURExpress® In Vitro Protein Synthesis Kit (NEB) following the manufacturer’s protocols. DNA templates for T7 polymerase-driven in vitro transcription were generated by PCR-amplification from wild-type (N16961) or various mutant (NtoxR.M13L; NtoxR.SDdown; NtoxR.G1,2,3; NtoxR.G3; NtoxR.G3,4; NtoxR.G4,5) cell lysates using KOD Hot Start DNA polymerase (Novagen®, MilliporeSigma), and primers toxR-T7.F1/toxR-T7.R1 (Table S2). The DNA templates were designed to comprise the T7 polymerase promoter consensus sequence fused to the 5’UTR and coding sequences of *toxR* with a T7 transcriptional terminator at the 3’ end. No native *toxR* promoter sequences were included in the template, and thus all transcription was driven from the T7 promoter. The PCR products were gel-isolated without the use of any DNA visualizing reagents and purified using the QIAquick Gel Extraction Kit (Qiagen), as per the manufacturer’s recommendations. The DNA template was used at a concentration of 250 nM per 25 μl reaction. CsrA was purified as described above and used at a concentration of 640 nM per 25 μl reaction. RNaseOUT™ Recombinant Ribonuclease Inhibitor (Thermo Fisher Scientific) was added to every reaction, as per the manufacturer’s instructions. Following incubation for 2 hours at 37°C, the PURExpress reactions were mixed with an equal volume of Laemmli sample buffer, and the *in vitro* protein synthesis products were analyzed by 12% SDS-PAGE gel electrophoresis, followed by Coomassie staining, as described above. ToxR synthesis in the presence or absence of CsrA was evaluated by immunodetection of ToxR, as described above.

## ACKNOWLEDGEMENTS

We gratefully acknowledge P. Julian Merville for technical assistance and Carolyn R. Fisher for critical reading of the manuscript. This work was supported by grants R01AI091957, R01AI16935, and R21AI124720 from the National Institute of Allergy and Infectious Diseases.

